# Human-specific expansion of 22q11.2 low copy repeats

**DOI:** 10.1101/2020.11.04.367920

**Authors:** Lisanne Vervoort, Nicolas Dierckxsens, Zjef Pereboom, Oronzo Capozzi, Mariano Rocchi, Tamim H. Shaikh, Joris R. Vermeesch

## Abstract

Segmental duplications or low copy repeats (LCRs) constitute complex regions interspersed in the human genome. They have contributed significantly to human evolution by stimulating neo- or sub-functionalization of duplicated transcripts. The 22q11.2 region carries eight LCRs (LCR22s). One of these LCR22s was recently reported to be hypervariable in the human population. It remains unknown whether this variability exists also in non-human primates. To assess the inter- and intra-species variability, we *de novo* assembled the region in non-human primates by a combination of optical mapping techniques. Orangutan carries three LCR22-mediated inversions of which one is the ancient haplotype since it is also present in macaque. Using fiber-FISH, lineage-specific differences in LCR22 composition were mapped. The smallest and likely ancient haplotype is present in the chimpanzee, bonobo and rhesus macaque. The absence of intra-species variation in chimpanzee indicates the LCR22-A expansion to be unique to the human population. Further, we demonstrate that LCR22-specific genes are expressed in both human and non-human primate neuronal cell lines and show expression of several primate LCR22 transcripts for the first time. The human-specificity of the expansions suggest an important role for the region in human evolution and adaptation.

**Author summary:** Low copy repeats or segmental duplications are DNA segments composed of various subunits which are duplicated across the genome. Due to the high level of sequence identity between these segments, homologous regions can misalign, resulting in reciprocal deletions and duplications, classified as genomic disorders. These regions are subject to structural variation in the human population. We recently detected extreme structural variation in one of the most complex segmental duplication regions of the human genome, the low copy repeats on chromosome 22 (LCR22s). Rearrangements between the LCR22s result in the 22q11.2 deletion/duplication syndrome, the most common human genomic disorder. However, it remains unknown whether this variability is human-specific. In this study, we investigated those LCR22s in several individuals of the different great apes and macaque. We show only the smallest haplotype is present without any intra-species variation in the *Pan* genus, our closest ancestors. Hence, LCR22 expansions are human-specific, suggesting a role of these LCR22s in human evolution and adaptation and hypothesize the region contributes to the 22q11.2 deletion syndrome inter-patient phenotypic variability.

## Introduction

Segmental duplications or low copy repeats (LCRs) constitute over 5% of the genome [1] and are complex patchworks of duplicated DNA fragments varying in length with over 90% sequence identity [2,3]. This high sequence homology has so far impeded the accurate mapping and assembly of these regions in the human reference genome [4,5]. Although it has become evident that assembly using short read sequencing is unable to resolve these complex regions, some LCRs are often too long and complex even for more recently developed long read technologies to resolve [4,6]. In addition, large structural variation amongst haplotypes complicates the assembly of these LCR containing regions [7]. As a consequence, LCRs remain poorly mapped and characterized, despite their functional importance in evolution and disease.

The impact of these LCRs on primate and human evolution is increasingly recognized [8,9]. It is estimated that the origin of the LCRs coincide with the divergence of New and Old World Monkeys, 35-40 million years ago [10]. However, a genomic duplication burst was observed in the great ape lineage, creating lineage-specific LCRs which are highly copy number variable [11]. These LCR-containing regions in other great ape reference genomes are also enriched for gaps, since they are subject to similar assembly difficulties as those encountered in the assembly of these regions in the human reference genome [12,13]. In humans, the 22q11.2 region contains a relatively higher proportion of LCRs compared with the rest of the genome. The origin of the human chromosome 22 LCRs (LCR22s) is concordant with the evolutionary timeline of LCRs in general. No duplicated orthologous LCR22 sequences are present in the mouse [14,15], and FISH mapping and sequencing experiments suggest lineage-specific LCR22 variation and mosaic structure in great apes [15-18]. However, since techniques to resolve the structure of the LCR22s were lacking, the great ape LCR22s have not been assembled and their composition and structure remain unresolved.

Due to the high level of sequence identity, homologous segments within LCRs can misalign during meiosis, via a mechanism known as non-allelic homologous recombination (NAHR), resulting in genomic rearrangements including deletions, duplications, and inversions [19]. The four most proximal LCR22 blocks are referred to as LCR22-A, -B, -C, and -D [20]. NAHR between these LCR22s underlies the formation of the recurrent deletions associated with the 22q11.2 deletion syndrome (22q11.2DS) (MIM: 188400/192430), the most common microdeletion disorder in humans [20] as well as the reciprocal duplications of this region often associated with abnormal phenotypes (MIM: 608363) [20].

We demonstrated hypervariability in the organization and the copy number of duplicons within LCR22s, especially LCR22-A [21]. By combining fiber-FISH and Bionano optical mapping we assembled the LCR22s *de novo* and uncovered over 30 haplotypes of LCR22-A, with alleles ranging in size from 250 kb to 2000 kb within 169 normal diploid individuals [21]. Pastor et al. recently expanded the LCR22-A catalogue by haplotyping the complete alleles of 30 22q11.2DS families [22]. To determine whether this extreme haplotype variability is human-specific, we set out to chart the inter- and intra-species variability of these LCR22s in non-human primates (S1 Table). The structures of the great apes, including five chimpanzees (*Pon troglodytes*), one bonobo (*Pon poniscus*), two gorillas (*Gorillo gorillo* and *Gorillo berengei groueri*), six orangutans (*Pongo pygmoeus* and *Ponglo obelii*), and one rhesus macaque (Old World Monkey, *Mococo molutto*) were analyzed by using an LCR22-specific fiber-FISH. To map the broader region, one representative of each species was analyzed by Bionano optical mapping. We demonstrate the non-human primate haplotypes to be less complex compared to humans. No intra-species variability similar to humans was observed suggesting that the hypervariability of the human LCR22-A haplotype is of recent origin.

## Results

### Conservation of the syntenic 22q11.2 locus

To assess whether the overall structure of the 22q11.2 region was conserved, the syntenic regions were investigated by Bionano optical mapping in non-human primate cell lines. Optical mapping allows the detection of structural variation, by imaging long fluorescently labeled DNA molecules (>150kb) followed by *de novo* assembly and local haplotyping [23]. Subsequently, the assembled alleles were compared to the human reference genome hg38 (Fig 1A). The resulting 22q11.2 syntenic assemblies were validated by fiber-FISH experiments using BAC (bacterial artificial chromosome) probes targeting the regions flanking the proximal LCR22s (schematic representation in Fig 1B, S2 Table). Due to the low mapping rate between the rhesus macaque sample and the human reference genome, the Bionano analysis in this non-human primate could not be performed and the composition (Fig 1G) is only based on fiber-FISH results.

**Fig 1.**
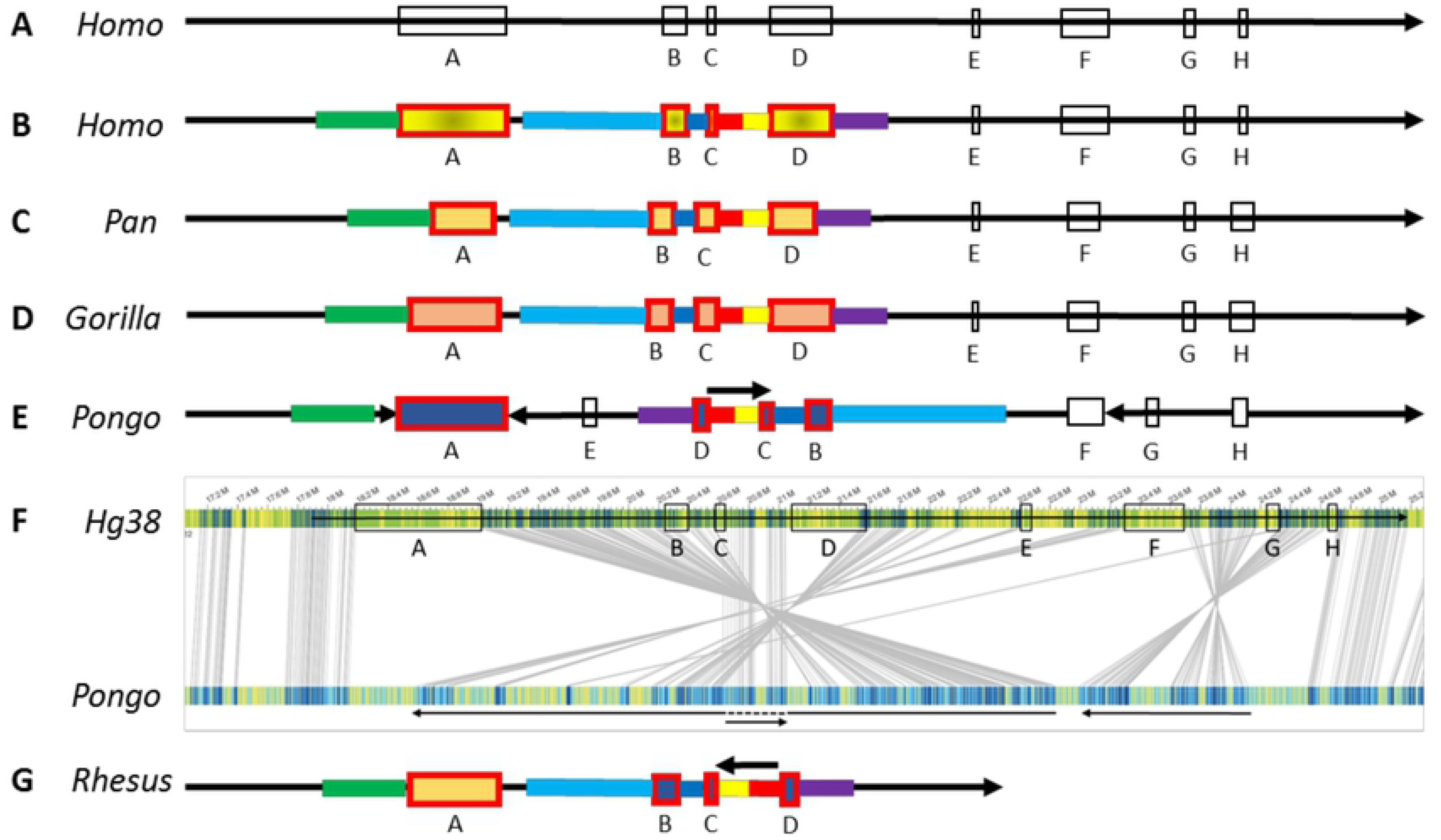
Composition of the 22q11.2 locus in human and non-human primates. Schematic representations of the 22q11.2 region, including LCR22-A through -H, based on Bionano optical mapping and fiber-FISH. As represented in (A) the human reference genome hg38, (B) human, (C) chimpanzee and bonobo, (D) gorilla, and (E) orangutan. (F) Bionano optical mapping results of orangutan compared to the human reference genome. The top bar represents the human hg38 reference genome with blocks indicating the LCR22s (corresponding to Figure 1A). The bottom bar represents the assembled haplotype for this orangutan. Grey lines between the maps indicate orthologous loci. Blue labels in the maps are aligned labels, and yellow labels unaligned. Arrows below depict rearrangements between the human and the orangutan genomes. (G) Schematic 22q11.2 representation of the macaque, only based on fiber-FISH results. Colored lines indicate the BAC probes used in the fiber-FISH experiments (S2 Table). Different sizes and colors of the LCR22 blocks indicate LCR22 differences in size and composition, respectively, based on fiber-FISH results. Cartoons are not to scale.

The order and organization of LCR22-A through -H in chimpanzee (Fig 1C, S1A Fig), bonobo (Fig 1C, S1B Fig), and gorilla (Fig 1D, S1C Fig) is identical to human. In contrast, three large rearrangements were observed in the syntenic 22q11.2 locus of the orangutan (Fig 1E-F). First, the region between LCR22-F and -H, including LCR22-G, is inverted. Second, an inversion is present between the LCR22-A and -F blocks. Third, the orientation between LCR22-C and -D is not inverted compared with the human reference. This could be interpreted as an extra inversion between LCR22-C and -D following the rearrangement between LCR22-A and -F. However, investigating this locus in the macaque by fiber-FISH uncovered the presence of this LCR22-C/D inversion, without the larger LCR22-A/F inversion (Fig 1G). Since we could not investigate the distal LCR22s, an inversion between these LCR22-F and -H cannot be excluded. Hence, despite the unstable nature of the LCR22s themselves, the structural organization between the LCR22 blocks is conserved between gorilla, chimpanzee, bonobo, and human. Inversions, typically flanked by LCRs, are present in the orangutan and rhesus macaque haplotype.

### Evolutionary analysis of LCR22-A

The current reference genomes of great apes, except for the chimpanzee, are enriched for sequence gaps within the loci orthologous to the LCR22s. As a consequence, it was not possible to fully rely on the reference sequences and alleles had to be *de novo* assembled. For this, an LCR22-specific fiber-FISH method was applied, which has proven its value to resolve these complex structures in humans (Fig 2A) [21]. Exact probe identities were checked by changing the fluorophores of color-identical probes (S2-5 Figs).

**Fig 2.**
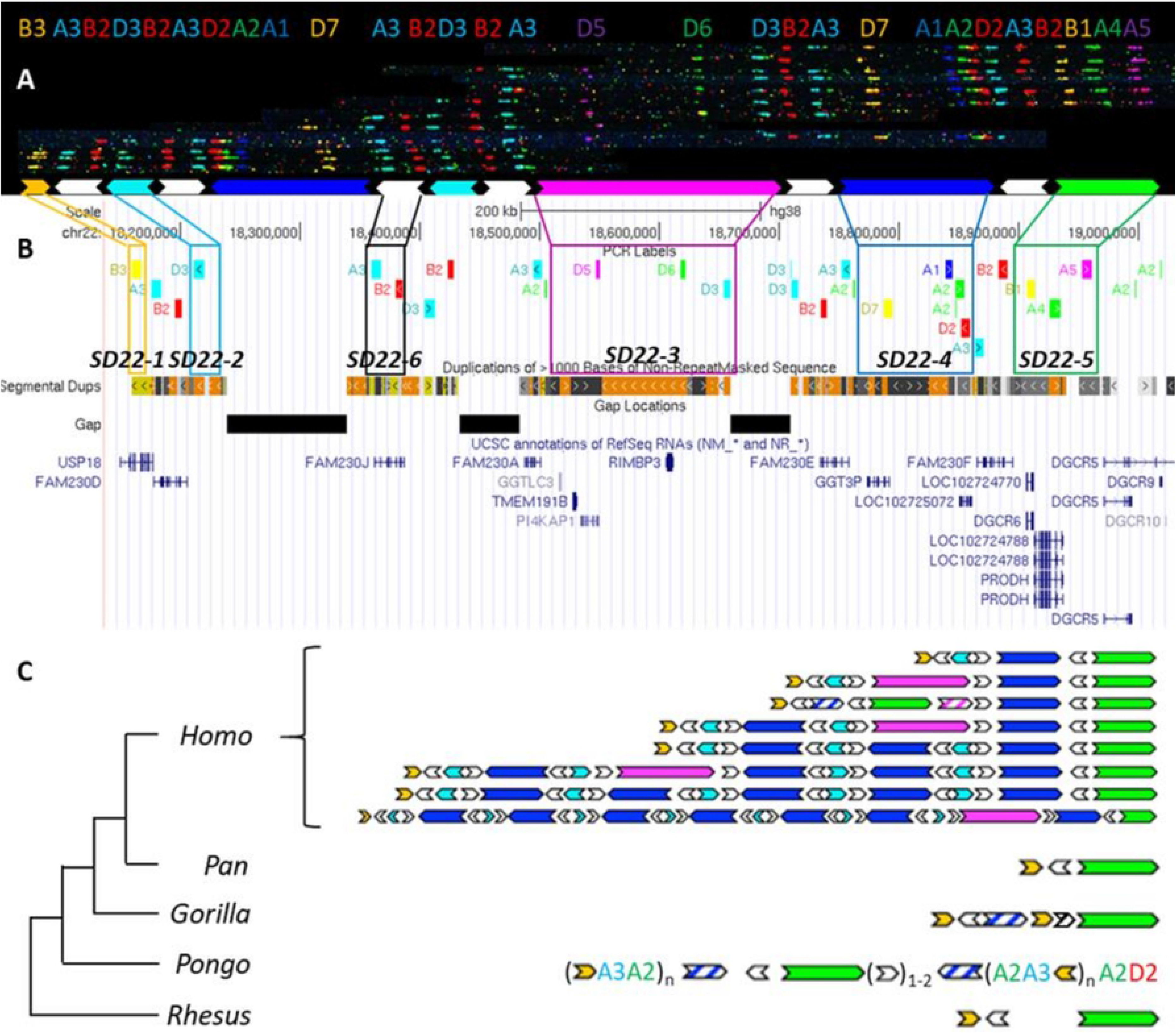
Human duplication structure and evolutionary analysis of LCR22-A. (A) *De novo* assembly of a LCR22-A haplotype based on matching colors and distances between the probes. SD22 duplicons are assigned to specific probe combinations. (B) UCSC Genome Browser hg38 reference screenshot, with tracks for fiber-FISH probe BLAT positions, segmental duplications, gaps, and RefSeq genes. Assigned duplicons in (A) are decomposed to their corresponding fiber-FISH probes in this reference screenshot. (C) Evolutionary tree representation of the observed LCR22-A haplotypes. Only a subset of assembled haplotypes are depicted for human, to emphasize the human hypervariability. Filled, colored arrows represent copies of duplicons, and hatched arrows represent partial copies of duplicons of the same color.

Based on the extensive variability observed in the overall size and duplicon content of human LCR22-A (Fig 2B-C), we wanted to determine whether similar variation exists in the other great apes and rhesus macaque. Towards this end, five chimpanzees, one bonobo, two gorillas, six orangutans, and one rhesus macaque were analyzed (S1 Table). In contrast to the human variability, no structural variation was observed in any of the ten alleles of LCR22-A observed in the chimpanzee samples (S6 Fig). In addition, both bonobo alleles also had the exact same composition as those in the chimpanzee. This haplotype (Fig 2C) is the smallest haplotype observed in human. However, this haplotype is rare in humans and only observed as a heterozygous allele in 5 of 169 human samples analyzed [21].

In the gorilla, the proximal and distal end are similar to the chimpanzee haplotype, except for a small insertion (Fig 2C). This is considered as a gorilla-specific insertion, since it is not present in the other non-human primate or human haplotypes. The same allele was observed in all four haplotypes of both gorilla cell lines. In addition to the large-scale rearrangements in the orangutan, we also observed major differences in the LCR22 compositions compared to the alleles of the other great apes (Fig 2C). First, the SD22-5 (green) duplicon, the distal delineating LCR22-A end in other great apes, is located in the middle of the allele, surrounded by SD22-6 duplicons. Second, tandem repeats, of probe compositions (indicated between brackets in Fig 2C) characterize the proximal and distal end of the allele. This characteristic is different from the interspersed mosaic nature of the LCR22s in humans. In addition, structural variation is observed within these repeats in the six orangutan samples (S3 Table). Thus, the haplotypes observed in the orangutan are very different from those observed in other great apes (Fig 2C). In contrast, the rhesus macaque haplotype is mostly identical to the small chimpanzee haplotype composition, except for an ~30kb insertion of unknown origin separating the SD22-5 and SD22-6 duplicons.

In order to validate these results, we correlated the fiber-FISH data with the corresponding chimpanzee reference genome. The human locus chr22:18,044,268-19,017,737 including the LCR22-A allele, can be traced to the chimpanzee locus chr22:2,635,159-2,386,886 in the most recent reference genome (Clint_PTRv2/panTro6/January 2018). The fiber-FISH probe order predicted from this sequence exactly matches the obtained fiber-FISH pattern. Hence, this extra independent chimpanzee allele confirms the presence of a single LCR22-A haplotype in chimpanzee.

In conclusion, due to the absence of LCR22-A structural variation in our closest ancestors, LCR22-A hypervariability can be considered as human-specific.

### Evolutionary analysis of LCR22-B/C/D

While LCR22-A is hypervariable in human genomes, LCR22-B and LCR22-C showed no variations, and only six different alleles were observed for LCR22-D (Fig 3A-D) [21]. To evaluate the evolution of these LCR22s and asses intra-species variation in non-human primates, we investigated the syntenic LCR22-B, -C, and -D haplotypes in great apes and rhesus macaque by fiber-FISH as well. Since LCR22-B and -C could be small and hard to distinguish above fiber-FISH noise, the probe set was supplemented with BAC probes flanking these LCR22s (S2 Table).

**Fig 3.**
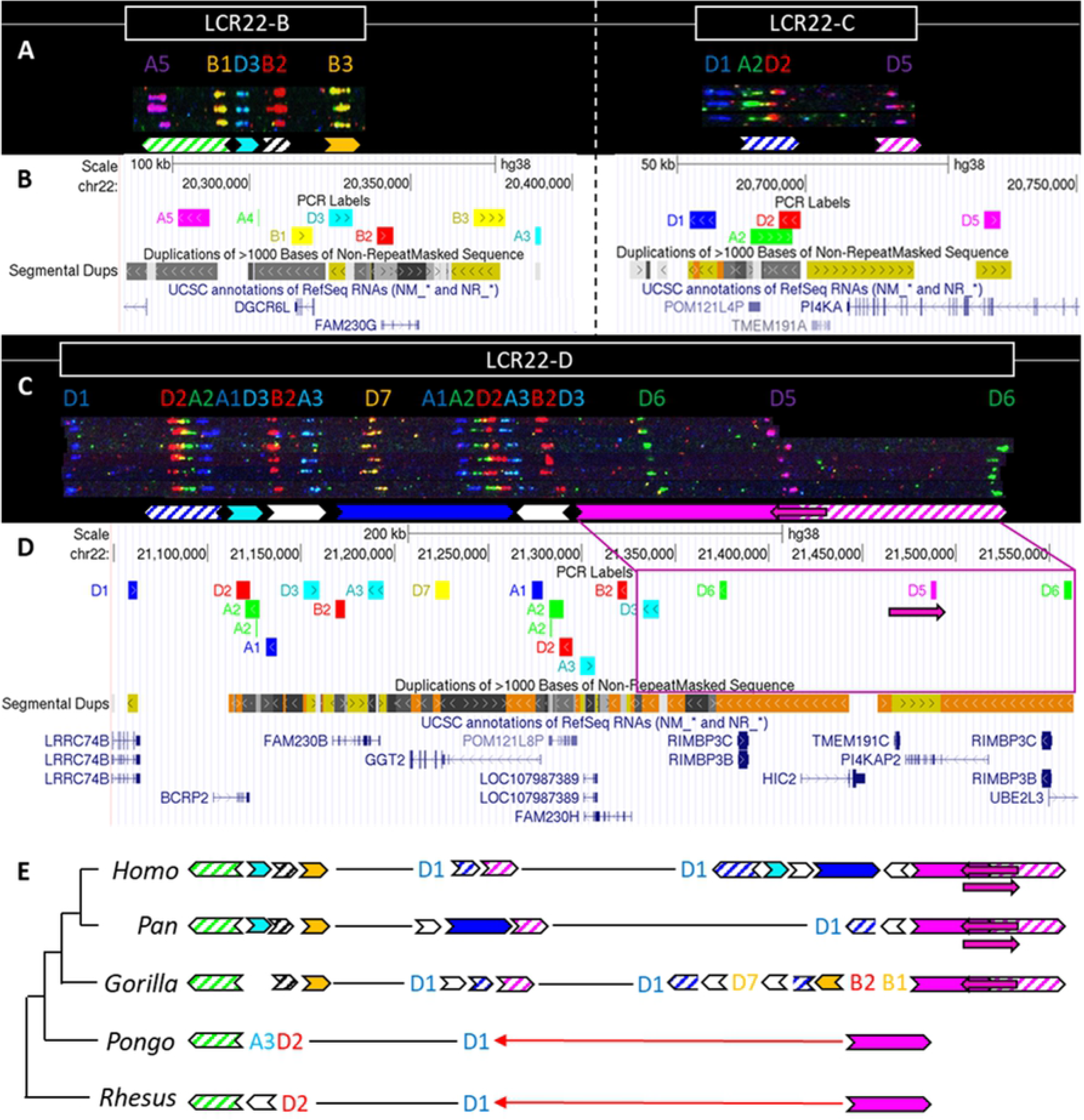
Human duplicon structure and evolutionary analysis of LCR22-B, -C, and -D. (A) *De novo* assembly of a LCR22-B (left) and LCR22-C (right) haplotype. SD22 duplicons are assigned to specific probe combinations, based on the probe composition in LCR22-A (Figure 2A). (B) UCSC Genome Browser hg38 reference screenshot of LCR22-B (left) and LCR22-C (right), with tracks for fiber-FISH probe BLAT positions, segmental duplications, and RefSeq genes. (C) *De novo* assembly of an LCR22-D haplotype based on matching colors and distances between the probes. SD22 duplicons are assigned to specific probe combinations. (D) UCSC Genome Browser hg38 reference screenshot, with tracks for fiber-FISH probe BLAT positions, segmental duplications, and RefSeq genes. The extended SD22-3 duplicon is decomposed to the corresponding fiber-FISH probes in the reference genome. (E) Evolutionary tree representation of the observed LCR22-B, -C, and -D haplotypes. Filled, colored arrows represent copies of duplicons, and hatched arrows represent partial copies of duplicons of the same color.

For LCR22-B, the chimpanzee and bonobo were identical to the human haplotype, while the gorilla haplotype was similar, with the deletion of one duplicon (SD22-2) (Fig 3E). In the orangutan, the distal part is substituted by two probes (A3-D2). An extra insertion between these two probes creates the haplotype of the rhesus macaque. LCR22-C carries lineage-specific insertions and deletions in the *Pon* and *Gorillo* genus, while in the orangutan and rhesus macaque it is reduced to only one probe (D1) (Fig 3E). The human LCR22-D haplotype is subjected to structural variation, mainly in the SD22-3 duplicon [21]. One variant, an internal inversion (indicated by the magenta arrow in Fig 3C-D), is present in 37% of the human haplotypes. The same variant was observed in a heterozygous state in two LCR22-D chimpanzee alleles (Fig 3E, S6 Fig), suggesting this variant precedes the split of the human lineage. The proximal start and distal end were conserved in Gorilla, with extra insertions compared to the human and *Pan* haplotype (Fig 3E). No structural variation was found at the distal end in these four investigated alleles. However, we predict that this structural variant is likely to be present in the gorilla population as well, since the composition is the same as in human, chimpanzee, and bonobo. The LCR22-D haplotype in orangutan and rhesus macaque is composed of only two probes (Fig 3E). To conclude, LCR22-B, -C, and -D haplotypes start to evolve towards their human structures in a common ancestor of *Gorilla, Pan* and *Homo*, based on the very short haplotypes found in orangutan and macaque.

### LCR22-specific transcript expression in human and non-human primates

According to the human reference transcriptome and the GTEx expression profiles [24], the LCR22s contain several expressed genes, pseudogenes, and long non-coding RNAs (Figs 2B, 3B, 3D). However, an expression study analyzing the LCR22 genes and their paralogs has not yet been accomplished in human nor in non-human primates. In addition, very few LCR22-specific genes have been annotated in the non-human primates. Based on the fiber-FISH composition of the non-human primate LCR22 alleles, we could predict the presence or absence of certain transcripts, since probes used in the fiber-FISH assays typically cover those genes. Hence, we set out to explore the conservation of the LCR22 specific genes and the expression pattern similarities with humans, by mining published primate transcriptome datasets. We explored gene expression in two publicly available brain transcriptome studies on human and non-human primates [25-27] (Table 1). The brain model was chosen since it is known that part of the human LCR22 transcripts are expressed in this tissue type, and genes within LCRs in general play a role in synaptogenesis, neuronal migration, and neocortical expansion in the human lineage [8,24].

**Table 1:**
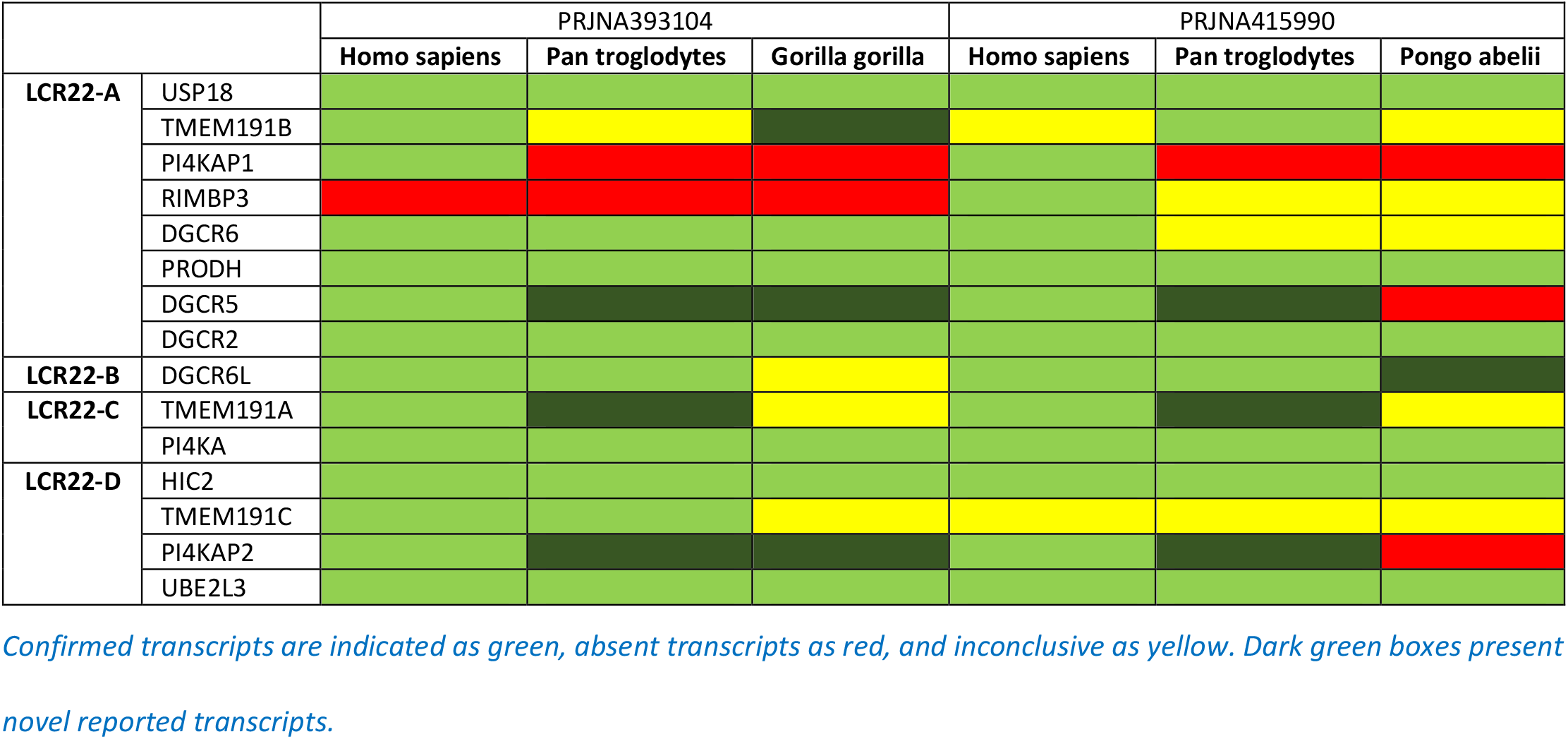
Transcriptome analysis of LCR22 genes for two publicly available transcriptome studies

We relied on two independent methods for the detection of transcripts: alignment and *de novo* assembly. Transcript assembly or alignment were seen as inconclusive when the coverage was below four reads or when paralogs could not be distinguished from each other. The latter was more frequently the case for non-human primates, as we lack a reference sequence of LCR22s and few orthologous transcripts have yet been annotated. Consequently, the *de novo* assembly has led to the discovery of several new transcripts for each of the non-human primates and a number of new splice forms of the *TMEM191* transcripts in the human genome. Moreover, *DGCR5* and *TMEM191A* are detected for the first time in non-human species (Table 1).

We attempted to distinguish between paralogs by adapting an assembly method originally developed for heteroplasmy detection in mitochondrial genomes [28]. Since this method needs sufficient coverage, we selected *PI4KA* and the two pseudogenes *PI4KAP1* and *PI4KAP2. PI4KA* was present in high coverage for all samples, while *PI4KAP1* was only found in human and *PI4KAP2* was only absent for orangutan (S1 Appendix). This pattern correlates with the fiber-FISH duplicons. *PI4KAP1* is located in SD22-3 in LCR22-A, which is unique to human (Fig 2). *PI4KAP2* is located in SD22-3 of LCR22-D, which is present in all great apes (Fig 3E). However, the SD22-3 duplicon in orangutan probably expresses the *PI4KA* gene, since the partial SD22-3 is absent in LCR22-C and the region is inverted. Therefore, absence of the *PI4KAP2* pseudogene in this species correlates with the absence of an extra SD22-3 duplicon in the fiber-FISH pattern. Although we were also able to identify some of the *TMEM191* paralogs with this method, low coverage, and the presence of multiple splice variants made it impossible to verify all paralogs in each study.

We looked at the expression of 39 LCR22 genes for two publicly available transcriptome studies (PRJNA393104 and PRJNA415990). Genes without distinct evidence of expression in any of the samples were excluded from Table 1 (a full list can be found in the S4 Table). For both transcriptome studies, there is clear evidence of LCR22 specific transcripts with a conserved expression pattern across both human and non-human primates (Table 1).

## Discussion

FISH mapping studies of metaphase chromosomes from great apes using 22q11.2 BAC probes and analysis of sequencing data had demonstrated the LCR22 expansion to precede the divergence of old and New world monkeys, and suggested species specific LCR22 variation had occurred during primate speciation [15-18]. However, the FISH studies were mainly focusing on interrogation of the copy number of a limited number of genic segments and sequencing analysis was inevitably interpreted against human reference genome 37 (hg19), carrying important inconsistencies compared to the most recent reference genome hg38. By *de novo* assembling the LCR22s using LCR22-specific probes in the fiber-FISH assay we resolved the haplotype composition in five chimpanzees, one bonobo, two gorillas, six orangutans and a macaque. This evolutionary analysis of the complex segmental duplications on chromosome 22 in different members of each species reveals that hypervariability of the LCR22-A allele is human-specific.

Human-specific expansions of LCR22s had introduced additional substrates for LCR22-mediated rearrangements which can result in genomic disorders associated with the 22q11.2 locus. As demonstrated by Demaerel et al. [21], the region of overlap between LCR22-A and LCR22-D is within a long stretch of homology encompassing SD22-4 flanked by SD22-6 on both sides, where recombination was shown to have taken place in case of an LCR22-A/D deletion. This locus is not present in any of the LCR22 blocks of the *Pan* genus. Pastor et al. [22] narrowed this region to SD22-6, the duplicon encompassing the *FAM230* gene member. Guo et al. [29] predicted the rearrangement breakpoint was located in the *BCR* (Breakpoint Cluster Region) locus, present in the distal part of SD22-4 (end of arrow). This locus was present twice in the *Pan* haplotype, once in LCR22-C and once in LCR22-D, but in opposite orientation preventing recombination leading to deletions and duplications. In the human lineage, the prevalence of both SD22-4 and SD22-6 increases in LCR22-A and LCR22-D. Hence, humanspecific expansion of the region likely increases the susceptibility of chromosome 22q11.2 to rearrangements, similar to observations made in other diseases resulting from LCR-mediated rearrangement [30].

The *Pan-Rhesus* LCR22-A haplotype is the smallest amongst the apes and was present in a homozygous way. Hence, this is likely the ancestral haplotype, with lineage-specific insertions and deletions. This ancestral haplotype is composed of three core duplicons (SD22-1, SD22-6, and SD22-5). Compared with most human haplotypes, three other core duplicons are missing (SD22-2, SD22-3, and SD22-4). These elements are present in respectively LCR22-B/D, LCR22-D, and LCR22-C of the *Pan* genus. Babcock et al. [31] presented a model of insertion of duplicons into LCR22-A combining homologous recombination in the absence of a crossover with non-homologous repair. The model was proposed for an interchromosomal recombination, but can be applied for intrachromosomal events as well. Following insertion in the LCR22-A block, allelic homologous recombination is a possible mechanism for the creation and expansion of new haplotypes. Since *Alu* elements are frequently delineating LCR blocks in general and on chromosome 22 specifically, they form a perfect substrate for this type of rearrangements [18,31,32].

This study provides the hitherto highest resolution map of the LCR22s across our closest evolutionary relatives, showing lineage-specific inversions, insertions, and deletions. Bionano optical mapping identifies three LCR22-mediated inversions in the orangutan lineage, and one in the rhesus macaque. A previous study focusing on the identification of inversion variants between human and primate genomes, observed the inversion between LCR22-C/D in the rhesus macaque, but was not able to identify any in the orangutan [33]. The extreme LCR22 amplification in gorilla, as described by Babcock et al. [17], was not identified in this study. It seems likely that some of the LCR22 duplicons are amplified at other regions in the gorilla genome. Since metaphase and interphase FISH studies have a lower level of resolution, the exact location of these amplifications is not known but some amplifications appear to be located at telomeric bands. Hence, they will not be identified by our LCR22 targeted fiber-FISH analysis.

It remains to be uncovered how this LCR22 variability influences the human phenotype and which elements in the regions are under selective pressure. Human-specific expansions were also observed in LCRs present on other chromosomes that are known to cause genomic disorders [34,35] and have been associated with human adaptation and evolution [8]. Gene duplications are a source for transcript innovation and expansion of the transcript diversity due to exon shuffling, novel splice variants, and fusion transcripts by the juxtaposition of duplicated subunits [36-38]. The human-specific *SRGAP2C* gene on chromosome 1 is an example of neofunctionalization [39]. The LCR-located gene, created by incomplete duplication, exerts an antagonistic effect on the ancestral *SRGAP2A* transcripts, resulting in human-specific neocortical changes [39,40]. Another example is the partial intrachromosomal duplication of *ARHGAP11A* (chromosome 15) leading to *ARHGAP11B*, which is associated with brain adaptations during evolution [41]. Hence, human-specific (incomplete) duplications of genic segments can render those genes into functional paralogs with possible innovating functions. These genes present evidence of positive selection and show a general increase in copy number in the human lineage [11].

The LCRs on chromosome 22 might be considered as an extreme source for expansion of the transcript catalogue. Several genes are present in the copy number variable duplicons of the LCR22s: *PI4KA* (SD22-3) and paralogs *PI4KAP1* and *PI4KAP2, RIMBP3* (SD22-3) and paralogs *RIMBP3B* and *RIMBP3C, FAM230* non-coding RNAs (SD22-6). First, most of these paralogs are not well characterized and classified (possibly incorrectly) as non-coding. We have clearly demonstrated expression of *PI4KA* (LCR22-C) and its non-processed pseudogenes *PI4KAP1* (LCR22-A) and *PI4KAP2* (LCR22-D). The expression is correlated with the presence or absence of the SD22-3 duplicons in the different species. Second, due to the high variability of these haplotypes in the human population, not every individual will have the same LCR22 genes or genic copy number. For example, due to this LCR22-A haplotype variability, the *PI4KAP1* pseudogene is not present in every human. Hence, the presence of specific paralogs and their possible functional importance might be underestimated. Transcriptome studies may help to unravel the role of these human-specific expansions. Short-read RNA-Seq datasets can be used to detect transcript expression (Table 1, S4 Table). Due to the duplicated nature of the LCR22s, paralogs share a high level of sequence identity. Therefore, short-read data are not always able to resolve the differences between transcripts arising from different paralogs. To unravel the predicted transcriptome complexity and the contribution of individual paralogs, long read full-length transcriptome analysis will be required. In addition, tools to obtain the full-length sequences of the LCR22s and map the paralog variability will be essential to fully comprehend the extent of sequence variation present. Our analysis focused on brain RNA-Seq datasets because of the importance of LCRs in the human brain development. However, absence of a transcript in the dataset does not automatically means that the gene is absent. For example, *FAM230* and *RIMBP3* paralogs are mainly expressed in testis [42-44]. LCR22-specific tissue transcriptome mapping or mining of the human cell atlas will be required to determine the full impact of the genes in those regions.

In summary, optical mapping of the LCRs on chromosome 22 unraveled lineage-specific differences between non-human primates and demonstrated the LCR22-A expansions and variability unique to the human population. It seems likely this expansion renders the region unstable and triggers NAHR resulting in the 22q11 deletions or duplications. To counter the paradox that LCR22 expansions reduce overall fitness, we hypothesize an important role for the region in human evolution and adaptation, previously described as the ‘core duplicon hypothesis’ [45-47]. Further research will be needed to unravel the functional importance of LCR22 expansion, including the role of paralog-specific transcripts.

## Materials and Methods

### Sample collection and cell culture

Four chimpanzee samples (*Pan troglodytes* 7, 8, 15, and 17), one gorilla cell line (*Gorilla gorilla* 1), and five orangutans (*Pongo pygmaeus* 6, 7, 8, 9, and 10) were kindly provided by Professor Mariano Rocchi (University of Bari, Italy). All these samples were Epstein-Barr virus (EBV) transfected cell lines and cultured according to standard protocols. One chimpanzee fibroblast cell line was purchased from the Coriell Cell Repository (AG 06939A). One gorilla fibroblast cell line (Gorilla Kaisi) was originally obtained from the Antwerp Zoo (Antwerp, Belgium). The orangutan fibroblast cell line and the rhesus macaque kidney cell line were obtained from the European Collection of Authenticated Cell Cultures (ECACC) Repository. One EBV cell line was established from bonobo Banya from the Planckendael Zoo (Mechelen, Belgium). More information on the samples is provided in S1 Table.

### Fiber-FISH

Long DNA fibers were extracted from the cultured cell lines using the FiberPrep^®^ DNA extraction kit (Genomic Vision). The slides were hybridized with the LCR22-specific customized probe set[21], supplemented with BAC probes targeting the unique regions between the LCR22s (S2 Table). Probes were labeled with digoxigenin-dUTP (Jena Bioscience), fluorescein-dUTP (Jena Bioscience), biotin-dUTP (Jena Bioscience), or combinations of these, using the BioPrime DNA Labeling System (Thermo Fisher Scientific). Indirect labeling with Alexa Fluor 647 IgG Fraction Monoclonal Mouse Anti-Digoxigenin (pseudocolored blue, Jackson Immunoresearch), Cy3 IgG Fraction Monoclonal Mouse Anti-Fluorescein (pseudocolored green, Jackson Immunoresearch), and BV480 Streptavidin (pseudocolored red, BD Biosciences) detected the primary labeled probes. The slides were scanned by an automated fluorescence microscope (Genomic Vision) at three excitation levels, corresponding to the three fluorophores. Images were automatically compiled by the system. The slides were visualized in FiberStudio (Genomic Vision) and manually inspected for regions of interest. Based on matching colors and distances between the probes, alleles were *de novo* assembled.

### Bionano optical mapping

High-molecular weight DNA from one chimpanzee (*Pon troglodytes* 15), one bonobo (Bonobo Banya), one gorilla (*Gorillo gorillo* 1), one orangutan (*Pongo pygmoeus* 8), and the rhesus macaque was extracted using the SP Blood & Cell Culture DNA Isolation kit (Bionano Genomics) and labeled using the DLS DNA labeling kit (DLE-1 labeling enzyme, Bionano Genomics). Samples were loaded onto Saphyr Chips G2.3 (Bionano Genomics), linearized, and visualized using the Saphyr Instrument (Bionano Genomics), according to the Saphyr System User Guide.

All analyses were performed in Bionano Access (Bionano Genomics). After general quality assessment via the Molecule Quality Report, a *de novo* assembly was performed against the most recent human reference genome hg38. Structural variants could be detected at the genome-wide level in the generated circos plot. The 22q11.2 region was visually inspected for structural rearrangements by zooming in to this region and comparing the compiled haplotypes with the hg38 reference.

### Transcriptome analysis of LCR22 genes

We selected two transcriptome studies, one across eight brain regions (PRJNA393104) [25], and one of neuronal differentiated induced pluripotent stem cells (PRJNA415990) [26,27]. The publicly available transcriptome datasets were downloaded from the European Nucleotide Archive: 32 datasets were from PRJNA393104 and 22 from PRJNA415990. A full list of accession numbers can be found in S4 Table. For each study, samples of the same individual were pooled together to generate a higher overall coverage of each transcript. The pooled FASTQ files were aligned to the human reference genome with BWA-MEM [48] and converted to BAM files with SAMtools [49]. The de novo assemblies were executed with NOVOLoci, a targeted assembler under development that was modified for transcriptome data. NOVOLoci needs a seed to initiate the assembly, therefore we prepared a short seed (100-250bp) for each of the 39 genes of the LCR22s. The resulting assemblies were aligned to the NCBI database with BLAST. Paralogs were identified and assembled separately with the heteroplasmy module [28] of NOVOPlasty [50]. We manually inspected the transcriptome alignments to the LCR22s and observed a large fraction of reads within introns, which also manifests in the *de novo* assemblies as additional intronic sequences at the end of some transcripts. As we did not observe genomic contamination, the presence of intronic sequences most likely originates from nascent RNA [51]. These nascent RNA sequences were removed from the *de novo* assemblies based on coverage difference and visual alignment.

## Supporting information captions

**S1 Fig. Bionano optical mapping of the 22q11.2 region in chimpanzee, bonobo, and gorilla.** Regional organization of the 22q11.2 locus in (A) chimpanzee, (B) bonobo, and (C) gorilla. De novo assembled non-human primate maps are compared to the human reference genome (hg38). The top bar represents the human hg38 reference genome with blocks indicating the LCR22s. The bottom bar represents the assembled non-human primate haplotype. Grey lines between the maps indicate orthologous signals between them. Blue labels in the maps are aligned labels, and yellow labels unaligned.

**S2 Fig. Exact probe composition of the LCR22 chimpanzee haplotypes.** To derive the exact probe composition of the chimpanzee haplotype, color-identical probes were differently labeled and hybridized to the slides. Changes of the pattern indicate the presence of the differently labeled probe. Red, cyan, and yellow probes were checked.

**S3 Fig. Exact probe composition of the LCR22 gorilla haplotypes.** To derive the exact probe composition of the gorilla haplotype, color-identical probes were differently labeled and hybridized to the slides. Changes of the pattern indicates that the presence of the differently labeled probe. Red, cyan, blue, and yellow probes were checked.

**S4 Fig. Exact probe composition of the LCR22-A and -B orangutan haplotypes.** To derive the exact probe composition of the orangutan haplotype, color-identical probes were differently labeled and hybridized to the slides. Changes of the pattern indicate the presence of the differently labeled probe. Red, cyan, blue, magenta, green, and yellow probes were checked. LCR22-C and -D were not included in the analysis, since they only consist of one and two probes, respectively. The probes are linked to unique BAC probes, predicting their composition.

**S5 Fig. Exact probe composition of the LCR22-A and -B rhesus macaque haplotypes.** To derive the exact probe composition of the rhesus macaque haplotype, color-identical probes were differently labeled and hybridized to the slides. Changes of the pattern indicates the presence of the differently labeled probe. Red, cyan, and yellow probes were checked. LCR22-C and -D are not included in the analysis, since they only consist of one and two probes, respectively. The probes are linked to unique BAC probes, predicting their composition.

**S6 Fig. Chimpanzee LCR22-A and -D haplotypes in investigated samples.** De novo assembled haplotypes for LCR22-A and LCR22-D in the six investigated samples. Two chimpanzees (*Pon troglodytes* 7 and 15) showed structural variation distal in the LCR22-D haplotype. A white line distinguishes the two haplotypes.

**S1 Appendix. Paralog analysis by heteroplasmy mode of NOVOPlasty.**

**S1 Table. Overview of non-human primate samples.**

**S2 Table. BAC probes targeting unique regions surrounding the LCR22s.**

**S3 Table. LCR22-A structural variation in the orangutan samples.**

**S4 Table. LCR22 specific transcript expression in human and non-human primates.**

